# EEG Oscillations in Guided Mindfulness versus Mind-Wandering in Young Adults: Effects of Auditory Task Instruction and Naturalistic Video

**DOI:** 10.64898/2026.08.28.747724

**Authors:** Daniel A. Brown, Kelsey E. Schultz, Finn Simmons, Carey Parker, Kevin Nute, Nicole C. Swann, Christina M. Karns

## Abstract

Alpha and theta band EEG oscillations have been implicated in states of mindfulness meditation. However, results are inconsistent and the influence of testing environment variables is not well-characterized. We used EEG to measure the amplitude of brain oscillations in a mindful-attention versus mind-wandering condition, in which guided auditory instructions were interleaved with periods of silence, with and without accompanying naturalistic video projections. We generated precise measures of alpha and theta in a sample of 20 young-adult non-expert meditators, by identifying each participant’s individual alpha peak frequency (IAF) from posterior EEG electrodes and using it to define four individualized frequency bands: two low alpha bands (in 2 Hz increments below the IAF), one upper alpha band (from IAF to 2 Hz above IAF), and one theta band (4 Hz - 6 Hz below IAF). We found that the mindfulness manipulation significantly increased power in alpha ranging between 2 Hz below to 2 Hz above IAF, including peak alpha amplitude, compared with mind-wandering. Meanwhile, central theta amplitude was larger when auditory instructions were on versus off, and naturalistic video did not reliably modulate the EEG effects of mindfulness. We conclude that the acute effects of mindfulness in non-expert meditators are most consistently observed as increases of posterior alpha-band activity, and that auditory task-instructions should be accounted for in studies of guided meditation. Observed alpha increases may reflect induced states of calm or relaxation induced by mindfulness practice, as proposed in previous studies. These results may apply to mindfulness training or biofeedback therapies.

## Introduction

Mindfulness meditation is an effective tool against many forms of distress (Goyal et al., 2014; Khoury et al., 2017), including anxiety (Bamber & Morpeth, 2019), stress (Pascoe et al., 2017), depression (Reangsing et al., 2021, 2022), pain (Jinich-Diamant et al., 2020), and other symptoms of mental health conditions (Vancampfort et al., 2021). An understanding of its neural underpinnings may identify critical pathways to inform future interventions for well-being (Brandmeyer & Delorme, 2020; Navarro Gil et al., 2018). Moreover, the synergy of meditation and nature exposure has shown early promise (Djernis et al., 2019; Lymeus et al., 2017), perhaps reflecting natural environments’ capacity to support mental health and emotion regulation (Anderson et al., 2018; Bell et al., 2026; Bratman et al., 2021). More experimental research is needed to understand the extent to which natural scenes influence neural correlates of mindfulness.

Mindfulness is the intentional regulation of attention toward present-moment experience, accompanied by an attitude of openness and non-judgment (Kabat-Zinn, 1982; Lutz et al., 2008). Within meditation, mindfulness is situated within focused-attention and open-monitoring frameworks, depending on whether attention is directed toward a specific object (e.g., the breath) or one’s ongoing experience (Lutz et al., 2008; Matko & Sedlmeier, 2019). In the present study, mindfulness was operationalized as a deliberate attentional modulation, with guided instructions interleaved with silence, to promote a sustained awareness of present-moment experience in novice meditators.

Although neuroscientific research on the brainwave impacts of mindfulness and meditation has grown, a consensus on the underlying mechanisms of mindfulness has not been reached. Increased alpha and theta power are the most frequently reported EEG findings across meditation (Cahn & Polich, 2006; Lagopoulos et al., 2009; Lee et al., 2018) including mindfulness (Lomas et al., 2015), but inconsistencies across studies remain (Cahn & Polich, 2006; Lee et al., 2018; Rodriguez-Larios et al., 2021), highlighting a need for improved research design and more sensitive methods (Lee et al., 2018; Lomas et al., 2015; Travis, 2020). More specific frequency bands, such as subdivisions above or below the alpha peak, have been proposed to isolate specific frequencies modulated by meditation practice (Travis, 2020). Since alpha peak frequency is known to vary across individuals, an adjustment for individualized alpha peak frequency (IAF) is warranted (Klimesch, 1999). By aligning comparable portions of the alpha band across participants, IAF-aligned methods increase signal-to-noise and reduce spectral “smearing” in group averages. As such, they may be useful to identify specific modulations that are missed by broadband analyses.

Finally, the simple presentation of auditory or visual stimuli in guided meditations can exert strong influences on brain signal and should be controlled for. Visual stimulation robustly modulates posterior alpha rhythms (Adrian & Matthews, 1934), auditory stimulation similarly alters ongoing oscillations (Makeig, 1993), and audiovisual stimulation has been shown to reduce alpha power during both auditory and video stimuli in both psychedelic and non-drug control states (Mediano et al., 2024). As such, stimulus-driven effects could confound subtle attentional manipulations, such as a guided mindfulness practice.

We designed a study to measure the impacts of three main variables on EEG oscillations during: (1) a guided mindfulness meditation compared to guided mind-wandering control condition, (2) with and without the projection of a large-scale naturalistic video, and (3) during audio-guided meditation periods and periods of silence. The purpose of this research was to a) describe how neural oscillations change with mindfulness meditation in novice meditators, b) investigate the potential synergistic effects of naturalistic imagery to enhance markers of meditation, and c) investigate the impacts of stimulus task design on EEG brain measurement. To optimally resolve meditation-related oscillation activity, we identified frequency bands precisely in individual participants relative to their individual alpha peak frequency (Klimesch, 1999).

We hypothesized that mindfulness would alter EEG power spectral density (PSD) in alpha and theta bands, and that naturalistic imagery and auditory instructions would influence neural responses in the mindfulness versus mind-wandering conditions.

## Methods

### Participants

We collected data from 20 right-handed university students between the ages of 18 and 35 (M age = 20.5, SD = 4.25). Participants self-reported gender identity, and the sample comprised 14 participants identifying as women and 6 identifying as men. Self-reported race and ethnicity were 18 White/Caucasian, 1 Black/White, and 1 Asian. Socioeconomic status was not collected. Participants were recruited through the university’s online human subjects pool. All participants signed a consent form approved by the university institutional review board for research ethics, and either received assignment credit in a course or were paid $15 per hour.

The original analysis plan was based on a fully within-subjects 2×2×2 ANOVA design. An *a priori* power analysis was conducted using G*Power (Faul et al., 2007). According to previous findings of alpha and theta power changes during mindfulness meditation (Lee et al., 2018; Lomas et al., 2015) we conservatively anticipated a medium effect size (Cohen’s f = 0.25), with an α-level of 0.05, power (1−β) of 0.80, and nonsphericity correction (ε) = 0.75. Given our initial planned within-subjects 2×2×2 ANOVA design, the power analysis yielded a minimum of 19 participants to detect a medium effect. Our final sample of N=20 exceeded this requirement, and is reflective of previously-published studies (Attar, 2025; Bell et al., 2026; Braboszcz et al., 2017; Katyal & Goldin, 2021; Lagopoulos et al., 2009; Megha et al., 2026; Rodriguez-Larios et al., 2021).

### Missing data and final analysis approach

Technical recording errors resulted in incomplete data in a few cases. Due to human error, the Mindfulness condition recording was not saved for two participants, and Mind-wandering without Video Projections was not saved for three additional participants. This left us with a sample of 20 for Mind-wandering with Video Projections, 17 for Mind-wandering without Video Projections, and 18 for Mindfulness (both with and without Video Projections). To accommodate incomplete repeated-measures data and retain all valid observations without imputing data, we instead analyzed the data using linear mixed-effects models using restricted maximum likelihood (REML), which incorporates all available observations without imputing missing values, under a missing at random assumption. The fixed-effects structure of the mixed-effects models corresponded to the original 2×2×2 design, and participant-level variability was modeled using random effects. Model based estimated marginal means were used for interpretation.

### Procedures

The experimental manipulations for this report occurred within the context of a larger study that included other motor and decision-making tasks, lasting approximately 1 hour with the order of tasks randomized. Activities of interest for this report encompassed 20 minutes and are summarized in Figure 1. Tasks were performed twice – once with a large-format video projection of droplets falling into still water (Naturalistic Video on) and once with a static gray square matched for size and luminance (Naturalistic Video off). The video of the movement of water droplets hitting standing water was created from architectural features designed to promote mindfulness through a natural sense of the passage of time (Nute & Chen, 2018). Visuals were projected onto a large 3.8 x 6.3-meter screen located in front of the participant using a short-throw projector, Epson PowerLite 535W WXGA 3LCD. Conditions were further randomized by condition blocks.

**Figure 1.**
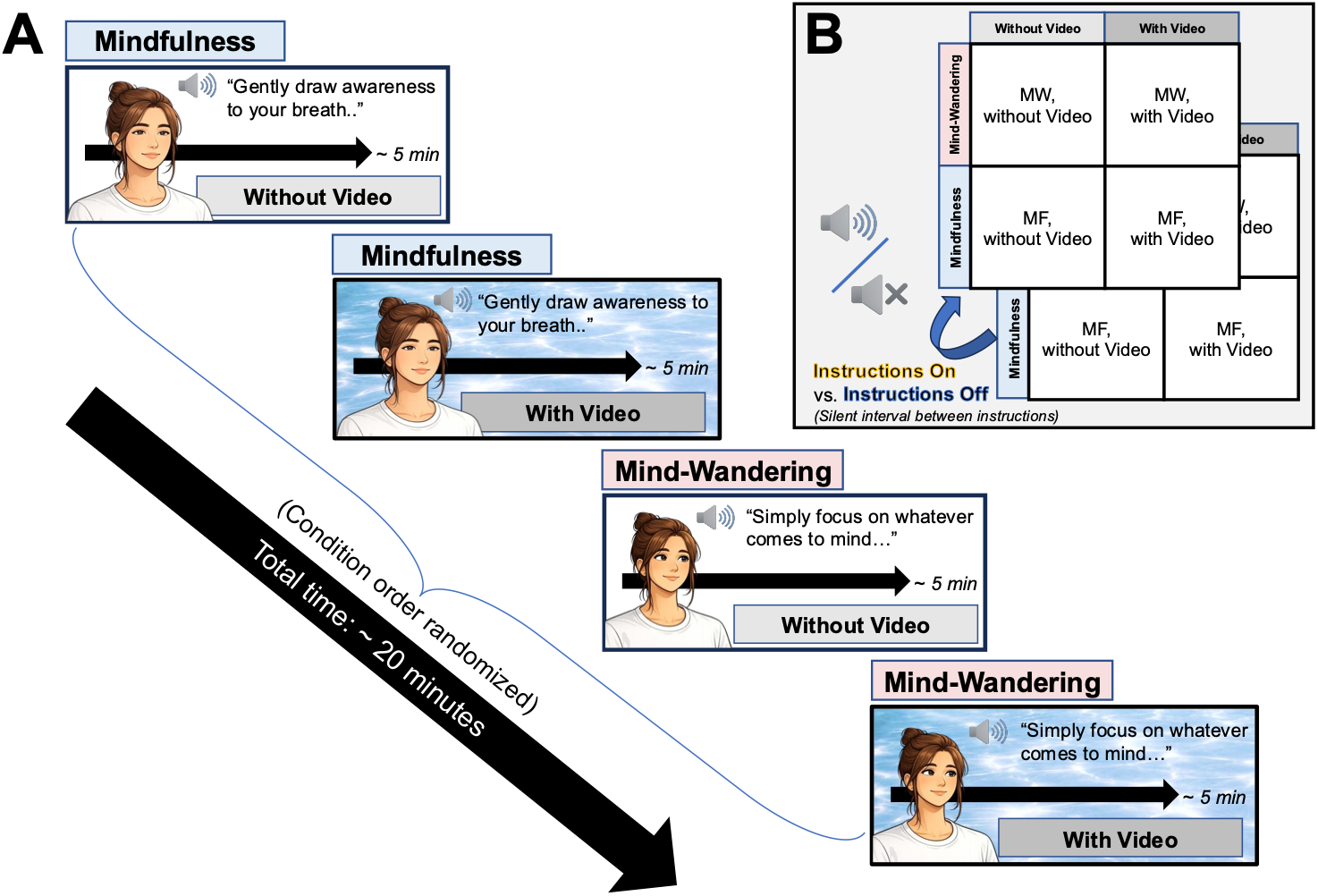
Visual depiction of the guided Mindfulness vs. Mind-wandering paradigm (A) and structure of the linear mixed-effects model design (B). During the naturalistic video projections, water refractions moved across the screen. Meanwhile, a gray square was used in the Without Video condition. Participants remained with their eyes open in all conditions. Key - MW: Mind-wandering. MF: Mindfulness.

In each condition block, subjects participated in a 5-minute guided session comprised of either a series of auditory Mindfulness instructions (e.g., “gently draw your awareness to your breath”), or a series of Mind-wandering instructions (e.g., “simply think about whatever comes to mind”). Instructions were recorded using a calm voice by a female narrator who was an experienced meditation guide. Within both Mindfulness and Mind-wandering guided sessions, auditory instructions lasted approximately 5 to 10 seconds, interleaved with 10 to 20 seconds of silence. Participants performed each unique condition once, experiencing four condition blocks in total. The onset and offset of each auditory instruction and silent period were additionally marked in the EEG recordings for data segmentation, creating a total of eight conditions for analysis. Participants completed all conditions with eyes open to preserve the primary manipulation of the visual environment. The structure of the task is depicted in Figure 1.

### EEG Recording

64-channel scalp EEG was recorded using an Active Two BioSemi system employing a CMS/DRL feedback loop during acquisition at a sampling frequency of 1024 Hz. Electrodes were positioned according to the standard 10-20 montage. External electrodes were positioned on the corner of the left eye, just above and below the left eye, and on the right and left mastoids. The electrodes around the eyes were used to measure ocular artifacts and aid with artifact correction, and the mastoid electrodes were recorded for re-referencing.

### Preprocessing

Signal preprocessing was conducted in EEGLAB (Delorme & Makeig, 2004). Channels identified as noisy via visual inspection were excluded from further analysis. A high-pass filter was then applied at 1 Hz to remove low-amplitude drifts, and the signal was re-referenced to the common average of the remaining EEG channels. Each file was then visually inspected for periods of high muscle noise or movement artifact, and periods of large paroxysmal artifact were removed.

Once the data had been pre-cleaned, independent component analysis (ICA) was used to identify and remove ocular artifacts using *runica* in EEGLAB (Jung et al., 2000), with data reduced to 30 dimensions using principal components analysis (PCA). ICA components were visually inspected and those corresponding to blinks and vertical or horizontal eye movements were removed. The cleaned EEG time series were then segmented into separate time series for each of the 2×2×2 conditions (Mindfulness/Mind-wandering, With/Without Video Projections, Auditory Instructions ON/OFF), for a total of 8 conditions.

#### EEG Analysis

##### Power Spectral Density (PSD)

Power spectral density (PSD) data analyses were conducted in MNE (Gramfort et al., 2013). For each participant and each of the 8 conditions, PSD from 1 to 45 Hz was calculated using the Welch method. Two-second sliding windows were used to encompass at least two full cycles of the lowest frequency of interest (1 Hz, the hi-pass filter value applied in preprocessing). Units of power were expressed in decibels (dB; 10 x log10 of μV^2^/Hz).

### Measurement of Frequency Bands Relative to Individualized Alpha

The precise frequency of ongoing oscillations corresponding to alpha is known to vary across individuals, with most individuals demonstrating a peak in their PSD between 7.5 Hz to 12.5 Hz (Klimesch, 1999). To account for this variation across individuals, we used a procedure described by (Klimesch, 1999) to define power bands for each subject based upon their individual alpha peak frequency (IAF). An example of applying this procedure to one participant is shown in Figure 2A.

**Figure 2.**
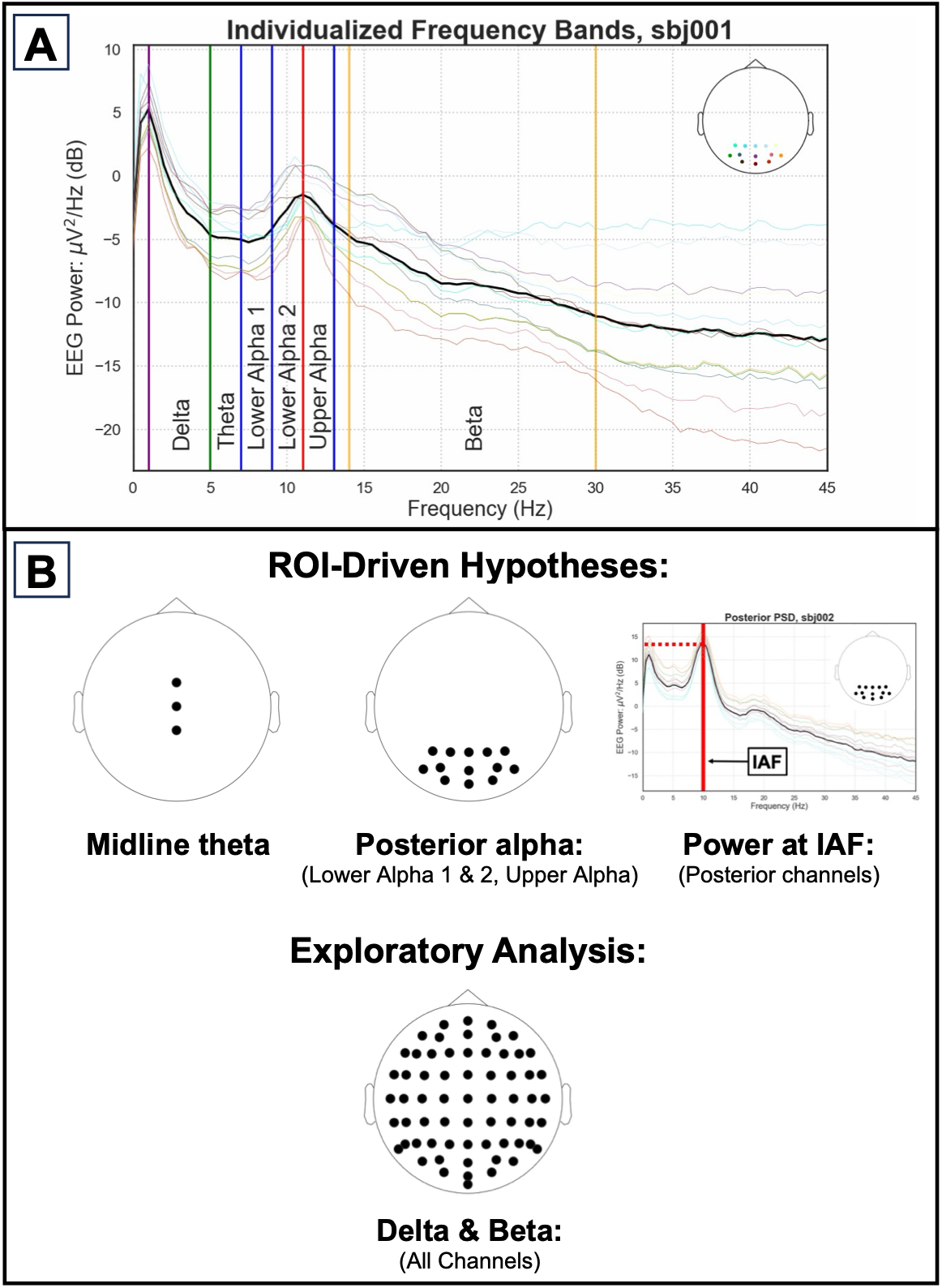
Individualized definition of frequency bands and linear mixed-effect model features. (A) Each participant’s individual alpha peak frequency (IAF; red line) was identified using the average power spectral density (PSD) curve from posterior scalp electrodes (bold black line) during both Mind-wandering and Mindfulness sessions combined. The IAF was determined as the peak value of the average posterior PSD, occurring within the range of 7 Hz and 13 Hz for each participant. Alpha and theta bands were then determined according to the IAF in 2 Hz increments. Delta was defined as 1 Hz to theta lower boundary. The beta lower boundary was determined by the larger of either 14 Hz or (IAF + 2 Hz), and the beta upward boundary was 30 Hz. (B) For alpha and theta, PSD power was extracted from regions of interest (ROIs) according to previous meditation findings (Cahn & Polich, 2006; Lagopoulos et al., 2009; Lee et al., 2018; Lomas et al., 2015). The power occurring at the individual alpha peak for each participant was extracted from posterior channels using the process identified in (A). In lieu of consistent previous findings to guide hypotheses, average delta and beta band power were calculated across all channels and included in linear mixed-effects models as exploratory analyses.

We used a region of interest (ROI) approach. Following (Klimesch, 1999), we calculated the PSD in each participant’s “posterior” EEG channels: [’P1’, ‘P3’, ‘PO3’, ‘P2’, ‘P4’, ‘PO4’, ‘Pz’, ‘POz’, ‘Oz’, ‘O1’, ‘O2’, ‘PO7’, ‘PO8’] during the entire 20 minute Mindfulness/Mind-wandering paradigm. These were selected because alpha peaks are most prominent in occipital and parietal channel locations (Adrian & Matthews, 1934; Klimesch, 1999). Resulting PSDs were then averaged to obtain a posterior average PSD (Figure 2A, bold black PSD curve). The IAF was automatically defined as the maximum power achieved between 7 and 13 Hz in the average posterior PSD. From prior datasets, we anticipated that some participants would not have a clear peak and would need alpha defined manually, either as an upward defection in the PSD slope or as the center of the 7-13 Hz range. Accordingly, IAF characterizations were visually confirmed for each participant, and the IAF was manually identified for one participant. Finally, we conducted follow-up analyses (Supplementary Figure 1) and confirmed that IAF characterization did not differ between posterior and frontal scalp electrodes.

The average power was extracted from IAF-centered frequency bands (Klimesch, 1999) as follows: upper alpha band (IAF to IAF + 2 Hz), lower alpha 2 band (IAF to IAF - 2 Hz), lower alpha 1 band (IAF - 2 Hz to IAF - 4 Hz), and theta band (IAF - 4 Hz to IAF - 6 Hz). Because the power spectral density is sampled in discrete 0.5 Hz bins, band edges were treated as inclusive at both ends; adjacent bands therefore share the single bin at each common boundary. Klimesch (1999) did not specify an IAF-based calculation for the delta band or beta band, so we defined them ourselves for exploratory analyses. Delta was defined as (1 Hz to IAF - 6 Hz). The lower boundary of the beta band was defined as the higher of (IAF + 2 Hz or 14 Hz), and the upper boundary of beta was 30 Hz (Figure 2A).

Once the boundaries of each frequency band were determined for each subject, the average PSD was extracted from scalp-based ROIs. In our hypothesis-driven regions, alpha was extracted from the same posterior electrodes used to define the IAF (see Figure 3 for group analyses), and theta was extracted from midline electrodes [’FCz’,’Cz’,’CPz’]. Although some mindfulness studies have reported changes in delta (Dunn et al., 1999) and beta (Ahani et al., 2014), these findings are not as prevalent or consistent as findings in alpha and theta (Lomas et al., 2015). Thus, our delta and beta power band measurements were extracted using all channels (Figure 2B), and analyses in these frequency bands were exploratory.

**Figure 3.**
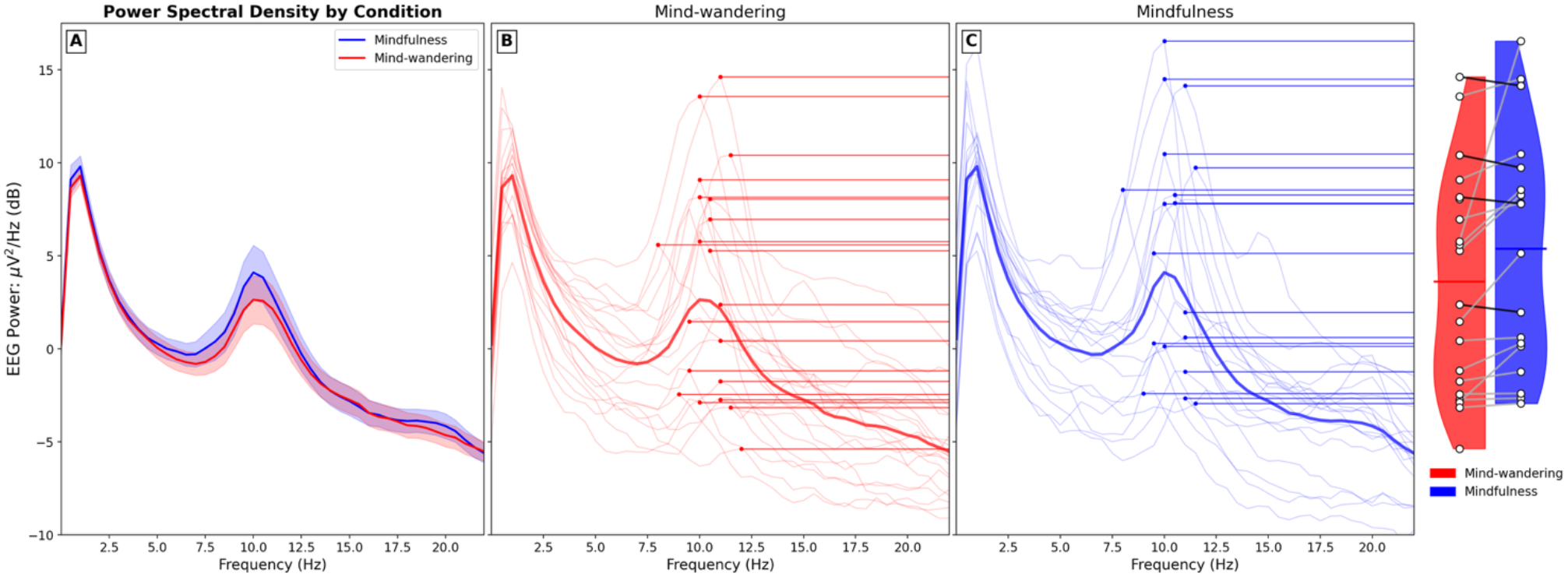
Power Spectral Density (PSD) by Mindfulness condition. (A) Mean Posterior PSDs: Unadjusted group mean PSD observed in posterior channels for Mindfulness vs. Mind-wandering conditions. This method exhibits “smearing," caused by averaging individual participants’ PSDs with varying peak alpha frequencies. (B) PSDs for individual subjects in the Mind-wandering condition reveal individual variability of alpha peaks; the amplitude observed at the individual alpha peak frequency (IAF; red dots) of each participant’s mean posterior-channel PSD (red lines; bolded = group average for condition). (C) PSDs for individual subjects in the Mindfulness condition. (D) Condition distribution comparison: Amplitudes extracted at the IAF were higher during guided Mindfulness vs. guided Mind-wandering (p<.001, p_FDR_ = .0063 in linear mixed-effects testing; see Figure 4 & Table 1 for full results). Lines link individual participants, with Mindfulness-related increases plotted as grey lines (n=14) and decreases plotted as black lines (n=4).

**Table 1.** Hypothesis-driven linear mixed-effects model results for EEG spectral outcomes. Linear mixed-effects models were fit separately for each EEG outcome with fixed effects of Mindfulness, Naturalistic Video, Auditory Instruction, and all two-way and three-way interactions, with participant included as a random intercept. Models were estimated using restricted maximum likelihood (REML), and inference was based on fixed effects. To control for multiple comparisons, main-effect tests for Mindfulness, Video, and Instruction across all seven EEG outcomes were adjusted using the Benjamini-Hochberg false discovery rate procedure (3 predictors x 7 outcomes) with p-values less than .001 treated conservatively as .0009. Nonsignificant interactions are omitted for brevity. Significance indicators reflect FDR-corrected p-values: * p_FDR_ < .05, ** p_FDR_ < .01.

| Outcome (ROI) | Effect | F(1, 119) | p<sup>raw</sup> | p<sup>FDR</sup> | Sig.<sup>FDR</sup> |
| --- | --- | --- | --- | --- | --- |
| **Theta (Central)** | Mindfulness | 3.65 | .058 | .245 |  |
|  | Video | 1.65 | .202 | .463 |  |
|  | Auditory Instructions | 7.22 | .008 | .043 | * |
| **Lower alpha 1 (Posterior)** | Mindfulness | 2.94 | .089 | .267 |  |
|  | Video | 0.39 | .533 | .733 |  |
|  | Auditory Instructions | 0.16 | .686 | .848 |  |
| **Lower alpha 2 (Posterior)** | Mindfulness | 16.00 | <.001 | <.0063 | ** |
|  | Video | 0.80 | .374 | .603 |  |
|  | Auditory Instructions | 1.11 | .294 | .560 |  |
| **IAF amplitude (Posterior)** | Mindfulness | 29.57 | <.001 | <.0063 | ** |
|  | Video | 1.57 | .213 | .463 |  |
|  | Auditory Instructions | 0.34 | .559 | .733 |  |
| **Upper alpha (Posterior)** | Mindfulness | 36.81 | <.001 | <.0063 | ** |
|  | Video | 3.23 | .075 | .261 |  |
|  | Auditory Instructions | 0.38 | .537 | .733 |  |

### Statistical Analysis: Linear Mixed-Effect Models

Statistical analyses were conducted in jamovi (v2.6) using the GAMLj module (Gallucci, 2023 & 2024; jamovi, 2024; Lüdecke, 2020; R Core Team, 2024). EEG power measures were analyzed using linear mixed-effects models, fit separately for each dependent variable, including: four alpha band power measures, each averaged across posterior electrodes (upper alpha, lower alpha 2, lower alpha 1, and the amplitude observed at the IAF), average theta band power observed in central electrodes, and two exploratory measures: average delta and beta power across all electrodes.

Because alpha power can vary with fatigue or task order, we first tested for correlations between block order and PSD measures to determine whether order should be included as a covariate. Because no significant relationships were detected, order was not included in the final models. Raw PSD data distributions within each condition were also inspected using box plots, which verified that no outlier values exceeded 1.5 of the interquartile range. Shapiro-Wilk tests confirmed normal distribution.

Each model included fixed effects of condition type (Mindfulness vs. Mind-wandering), naturalistic video projections (ON vs. OFF), auditory instruction (ON vs. OFF), and all two-way and three-way interactions. Participant was included as a random intercept to account for repeated measurements within individuals. Models were estimated using restricted maximum likelihood (REML), allowing all available observations with valid data for a given dependent variable to contribute to model estimation. Fixed effects were evaluated with Sattherthwaite approximations of degrees of freedom, and estimated marginal means were used to interpret significant effects and interactions.

To control for multiple comparisons, p-values for the main effects of Mindfulness, Naturalistic Video, and Instruction were adjusted across the seven EEG outcomes, including the exploratory delta and beta bands, using the Benjamini–Hochberg false-discovery-rate procedure (FDR; 3 predictors and 7 dependent variables for 21 tests total). P-values less than .001 were treated conservatively as p = .0009 in false-discovery-rate calculations. Linear mixed-effects findings appear in Figure 4 and Tables 1-2. Interactions were corrected separately from main effects, as they represent conceptually distinct tests: the Benjamini–Hochberg procedure was applied across the 28 interaction tests (four interaction terms × seven outcomes) as their own family. We planned to conduct post-hoc follow-up tests if any two- or three-way interaction was identified as significant; no interaction survived correction, and the simple effects reported in Supplementary Table 1 are therefore exploratory.

**Figure 4.**
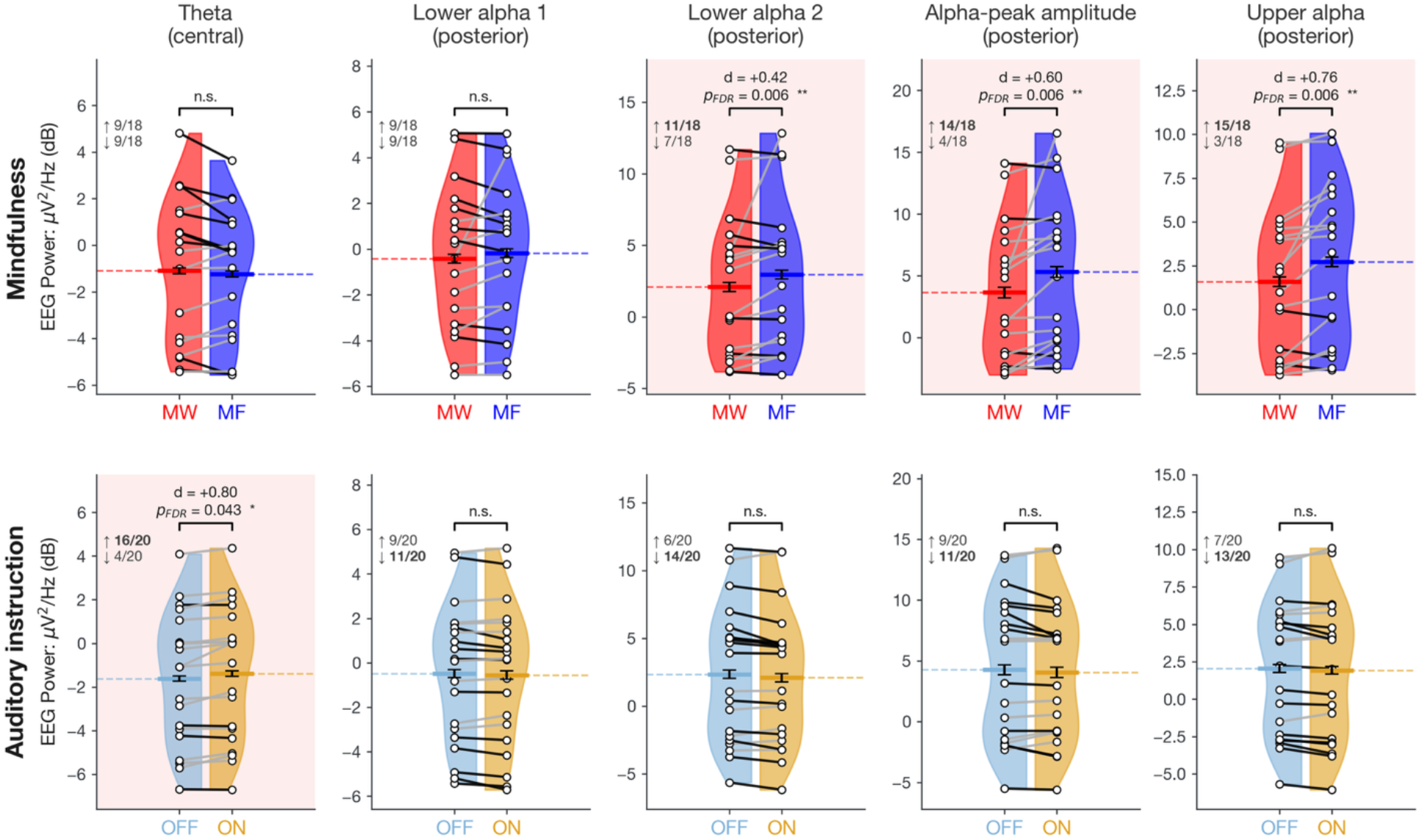
Hypothesis-driven linear mixed-effects outcomes for Mindfulness and Auditory Instruction contrasts. Columns show the five hypothesis-driven spectral outcomes; rows show the two contrasts that produced significant effects (Naturalistic Video yielded no significant contrasts). Panel shading in peach marks significant panels after FDR. Half-violins give the distribution across participants; dots mark individual participants, grey connecting lines indicate left-to-right increases, black lines correspond to decreases. Inset text shows proportion of subjects expressing increases or decreases, with winning direction bolded. Colored dotted lines indicate estimated marginal means, black error bars represent within-participant 95% confidence intervals from linear mixed-effects models. Effect sizes are within-participant Cohen’s dz; p-values are Benjamini-Hochberg corrected across the 21 main-effect tests reported in Table 1 and Table 2 (** p < .01, * p < .05).

**Table 2.**
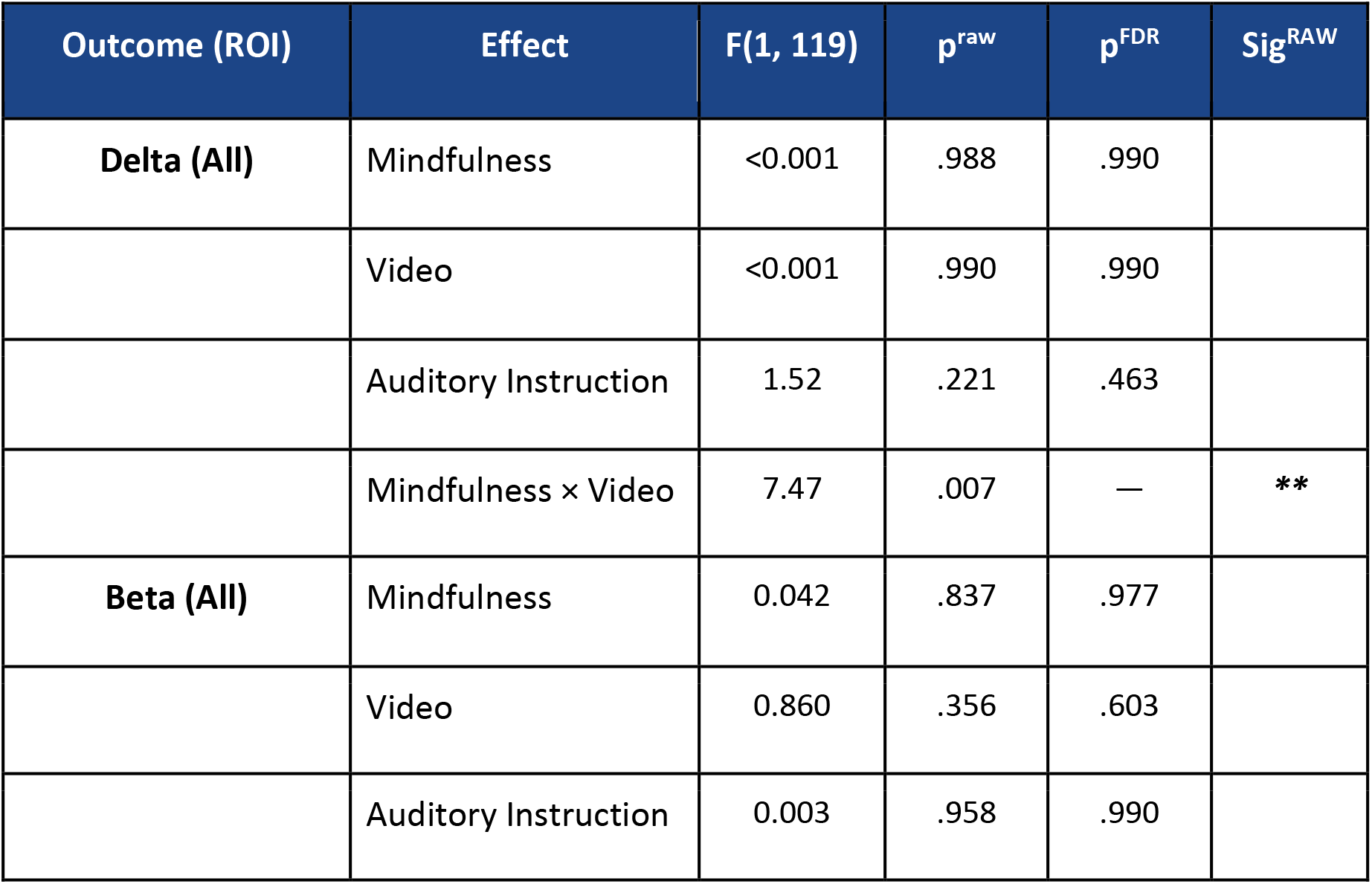
Exploratory analyses: Linear mixed-effects model results for EEG spectral outcomes. Exploratory analyses were conducted for delta and beta power using the same model structure described in Table 1. Main-effect p-values were included in the same 21-test FDR correction family as the hypothesis-driven outcomes. The Mindfulness × Video interaction for delta was excluded from the FDR family; its significance indicator reflects the raw p-value.

### Sensitivity Check: Fixed-band version of Linear Mixed-Effects Models

To demonstrate the statistical impact of employing IAF-aligned frequency bands, we re-ran the linear mixed-effects models with bands anchored to the group-mean IAF rather than each participant’s peak. Identical frequency band widths and FDR correction were applied as the main linear mixed-effects models. Findings from this analysis appear in Supplementary Figure 2, along with the main IAF-adjusted analysis for comparison.

### Cluster Analysis

Finally, we conducted an exploratory analysis of Mindfulness vs. Mind-wandering using cluster permutation testing across channel location and frequency (Figure 5). The goal of this test was to validate our hypothesis-driven findings using a data-driven approach, and investigate whether electrodes and frequencies outside of our initial hypotheses were affected by the manipulation. This test was run twice: once on IAF-adjusted spectra in which each participant’s spectrum was aligned to their own alpha peak, and once on non-adjusted spectra. Running both allowed us to directly quantify how individualized alignment contributes to model sensitivity.

**Figure 5.**
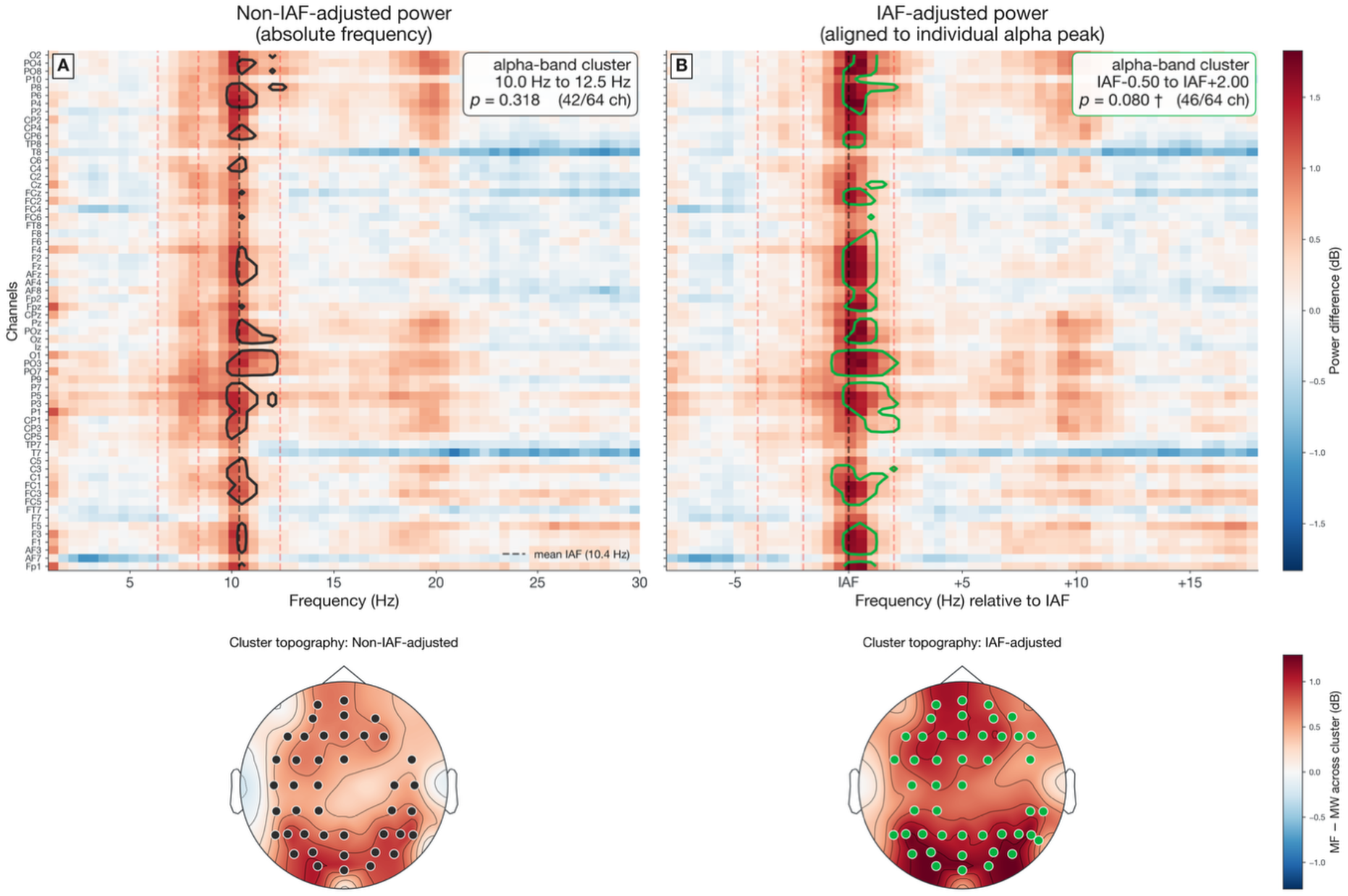
Individualized alpha frequency alignment improves sensitivity of data-driven cluster permutation testing. In n=18 participants contributing both conditions, IAF-aligned spatio-spectral testing of the Mindfulness - Mind-wandering contrast yielded trend-level alpha power findings, while non-adjusted methods did not approach significance. Top row: (A) In cluster permutation testing using non-adjusted spectra, the strongest cluster spanned 10.0–12.5 Hz, included 42 of 64 channels, and yielded a cluster-corrected p = .318. (B) When repeated with spectra aligned to each participant’s alpha peak, the corresponding cluster spanned (IAF – 0.50 Hz) to (IAF + 2.00 Hz), included 46 of 64 channels, and yielded a cluster-corrected p = .080 († denotes a trend-level cluster, p < .10). Dashed red lines in (B) mark (IAF - 4 Hz), (IAF - 2 Hz), IAF, and (IAF + 2 Hz), for comparison with the lower and upper alpha bands used in the linear mixed-effects models; in (A) these are positioned relative to the group-mean IAF (10.4 Hz, dashed black line). Bottom row: topographic shading indicating scalp distribution of the Mindfulness - Mind-wandering difference, averaged across the frequency range of the alpha-band cluster in each test. Dots mark the channels belonging to the identified clusters above.

Cluster permutation testing was conducted in MNE. Preparation for the analysis involved the following steps: In the IAF-shifted model, each participants’ PSD values were shifted according to their individual alpha peak frequency by subtracting the IAF (defined via posterior channels) from each PSD’s x-values, re-aligning every PSD to their posterior IAF. IAF-Aligned PSDs were then extracted for all channels in all participants for both the Mindfulness and Mind-wandering conditions, resulting in a matrix of 18 participants x 53 frequencies x 64 channels for each condition. Two participants were excluded due to missing data. The 53 frequencies ranged in equal 0.5 Hz steps from (IAF – 8 Hz) to (IAF + 18 Hz), matching the native frequency resolution of the Welch PSD so that the IAF-adjusted and non-adjusted tests search grids of the same resolution and clusters are directly comparable. The lower bound was set by the participant with the lowest IAF (8.0 Hz), below which not every participant has data after alignment. The upper bound of (IAF + 18 Hz) was chosen such that the range did not extend beyond beta (30 Hz) for any participant.

The Mindfulness and Mind-wandering matrices were then compared via cluster permutation testing in MNE (Gramfort et al., 2013) to identify clusters of adjacent channels/frequencies implicated in the comparison. Testing includes: a) performing 2-tailed within-subjects t-tests at each of the 64 × 53 spatio-spectral locations (64 channels × 53 frequency bins, spanning IAF − 8 Hz to IAF + 18 Hz at 0.5 Hz resolution), b) determining and applying a cluster-forming threshold at the 95% percentile of the t-value distribution, c) using a channel adjacency matrix to link together “islands” of spatial and spectral neighbors that exhibit significant comparisons, and d) performing 10,000 sign-flip permutations of the within-participant difference maps to form a null distribution against which cluster significance is assessed. This framework controls the family-wise error rate across the full channel x frequency search space, thereby providing correction for multiple comparisons while leveraging spatio-spectral clustering to maintain sensitivity. Family-wise error is controlled within each test but not across tests: the cluster analyses reported (IAF-aligned and non-aligned) are presented as one primary analysis (IAF-aligned; a replication of our main LME analysis) with a sensitivity variant.

## Results

### Linear Mixed-Effects Model Analyses: Main effects of Mindfulness Meditation

As a primary finding, for the main effect of Mindfulness vs. Mind-wandering, linear mixed-effects models revealed that Mindfulness increased multiple measures of posterior alpha-band activity. Per-participant and group-level outcomes for each model are reported in Table 1 and shown in Figure 4.

Upper alpha: Upper alpha power (0-2 Hz above IAF) showed a robust main effect of condition (Mindfulness vs. Mind-wandering), F(1, 119) = 36.81, p < .001 (p_FDR_ < .0063), with higher estimated marginal mean power during Mindfulness (M ≈ 2.60, SE ≈ 1.03) than Mind-wandering (M ≈ 1.45, SE ≈ 1.03). Mindfulness-related upper alpha increases were consistent across the Naturalistic Video and Auditory Instruction conditions, with no significant interaction effects, all raw p-values> .29. No other upper alpha main effects were significant after FDR correction. Model-estimated means and contrasts for interpreted fixed effects are reported in Supplementary Table 1.

Lower alpha 2: Posterior lower alpha 2 power (2 Hz below - IAF) showed a similar main effect of Mindfulness, F(1, 119) = 16.00, p < .001 (p_FDR_ < .0063), with higher power during Mindfulness than Mind-wandering. No other main effects or interactions in this frequency band were significant, all raw p-values ≥ .29.

Lower alpha 1: Posterior lower alpha 1 power (4 Hz below - 2 Hz below IAF) did not differ significantly between Mindfulness and Mind-wandering, F(1, 119) = 2.94, p = .089 (p_FDR_ = .267), and showed no significant main effects of Naturalistic Video or Auditory Instruction or significant interactions, all raw p-values ≥ .36.

IAF amplitude: Posterior power at the IAF frequency also showed a strong main effect of meditation, F(1, 119) = 29.57, p < .001 (p_FDR_ < .0063), with higher estimated marginal mean amplitude during Mindfulness than Mind-wandering. No other main effects or interactions at this frequency were significant, all raw p-values ≥ .21.

Other Frequency Bands: Theta and lower alpha 1 only showed trend-level main effects of Mindfulness at the uncorrected level (raw p-values = .058 and .089, respectively, p_FDR_ ≥ .24). Delta and beta bands showed no evidence of any condition-related differences (raw p-values ≥ .84, p_FDR_ ≥ .98).

### Auditory Instruction

Central theta power showed a significant main effect of Auditory Instruction, F(1, 119) = 7.22, p = .008 (p_FDR_ = .043), with higher estimated marginal mean power when instructions were ON than OFF. No other EEG frequency bands showed significant effects of Auditory Instruction after FDR correction (all p-values ≥ .20).

### Naturalistic Videos

There were no significant effects of Naturalistic Video projections at any frequency. A non-significant trend for posterior upper alpha power reflected slightly higher power during the no-video compared to video condition, which did not survive correction for multiple comparisons and should not be overinterpreted, F(1, 119) = 3.23, p = .075, (p_FDR_ = .261). All other frequency bands showed no evidence of even trend-level main effects of naturalistic video imagery (all p-values ≥ .20).

### Interactions

For delta power, an exploratory measure, an uncorrected Mindfulness × Naturalistic Video interaction was observed, F(1, 119) = 7.47, p = .007 (Table 2), indicating that the effect of Mindfulness may differ as a function of the Naturalistic Video condition. Applying Benjamini-Hochberg correction across the family of 28 interaction tests (four interaction terms × seven outcomes), this effect did not survive (q = .203), and no other interaction approached significance in any frequency band (all raw p-values ≥ .17). As this interaction was not part of our primary hypotheses and does not survive correction, we do not interpret it further.

### Fixed Bands variant

In mixed linear-effects models using frequency bands determined by group mean IAF rather than individual IAF, the Mindfulness effect on upper alpha was more than halved F(1, 119) = 36.81 vs. 16.68, and the estimated difference in this band was reduced from 1.14 dB to 0.71 dB (Supplementary Figure 2). A significant Mindfulness effect appeared in a band 2 - 4Hz below the mean IAF, corresponding to lower alpha 1, F(1, 119) = 11.01, p_FDR_ = .006, that individualized bands did not support, F(1, 119) = 2.94, p_FDR_ = .267. The alpha-peak amplitude measure was weakened under fixed anchoring, F(1, 119) = 23.54 versus 29.57.

### Summary

After FDR correction across all 21 main-effect tests, the Mindfulness effects on lower alpha 2, upper alpha, and IAF amplitude remained significant, as did the Auditory Instruction effect on central theta. No delta or beta main effects were significant after FDR correction. Trend-level uncorrected effects of Mindfulness on theta and lower alpha 1, and Naturalistic Video on upper alpha, did not survive correction.

### Exploratory Cluster Permutation Testing

We completed cluster permutation testing as a complement to our main analysis, to further investigate the frequencies and channels affected by the Mindfulness manipulation. The goals of this test were to validate our hypothesis-driven linear mixed effects model results in a data-driven manner, explore whether regions and frequencies outside our initial hypotheses were affected, and observe the statistical impacts of adopting individualized alpha-peak frequency binning.

Neither version of this exploratory analysis reached significance, but they differed in an informative way. Aligning spectra to each participant’s own alpha peak greatly reduced cluster-level p-values, and cluster testing recapitulated our main analysis findings that group differences were strongest at and just above the IAF (Figure 5). With IAF adjustment, the strongest of 11 clusters spanned (IAF - 0.50 Hz) to (IAF + 2.00 Hz) across 46 of 64 channels (p = .080), with its maximum at bilateral occipital sites (O1, PO8, PO3, PO7; Figure 5B). Without IAF adjustment, the strongest of 23 clusters occupied a comparable frequency range and a nearly identical scalp distribution (10.0 - 12.5 Hz across 42 of 64 channels, maximum effects at O1, PO3, PO8 and Oz), but the contrast was far weaker (p = .318). In sum, the implicated frequency range and topography were similar with and without alignment, but the effect was detected far more strongly once each participant’s spectrum was referenced to their own alpha peak – signaling the primary benefit of individualized alignment was increased sensitivity, rather than spectral or spatial specificity.

The cluster identified by IAF-adjusted testing included a wider set of EEG channels than our a priori posterior region, suggesting that Mindfulness-related alpha increases may extend beyond the channels we originally hypothesized. However, the largest effects remained within our hypothesized posterior ROI, and importantly the extended clusters did not reach statistical significance. Finally, a separate sensitivity analysis restricting both tests to the silent (Auditory Instructions Off) periods produced the same pattern (IAF-adjusted p = .096; Without IAF adjustment p = .293), indicating that our observed main effects were not carried by the instruction periods. In every version of these data-driven tests, including both IAF-adjusted and non-adjusted tests, the only other cluster of appreciable size fell in the beta range (approximately 18–21 Hz, equivalently IAF + 8.5 Hz to IAF + 11.5 Hz) but came nowhere near significance (all p-values ≥ .26).

## Discussion

In an era of endemic stress, mindfulness meditation has been linked to emotional, mental, and health-related benefits. Understanding its neural mechanisms may provide valuable insights for effective therapeutic interventions. The aim of this experimental study was to assess the impact of guided mindfulness meditation on EEG oscillations in novice meditators, with and without the presence of naturalistic visual stimulation and auditory instructions. The mindfulness condition increased measures of alpha defined in individual participants (posterior upper-alpha power, posterior lower-alpha 2 power, and power at the posterior individual alpha peak frequency). Auditory instructions also significantly modulated central theta power, with greater theta amplitude during interleaved instruction periods than during silent periods, which could reflect entrainment of theta to the speech envelope (Ding & Simon, 2014), but this did not explain the effect of mindfulness. In contrast, we did not find convincing evidence that naturalistic video stimulation significantly affected the EEG impacts of meditation. As a validation of our hypothesis-driven findings, we also utilized data-driven cluster permutation testing to investigate the specific frequencies most impacted by our Mindfulness manipulation. This analysis confirmed that frequencies around and above the IAF were most impacted by Mindfulness, but the method was less powerful. Across our data-driven and hypothesis-driven testing, we found confirmatory evidence that using individually-adjusted alpha-peak values in analysis greatly improved the sensitivity of our statistical models. Taken as a whole, we found that alpha power increases with guided mindfulness relative to guided mind-wandering, in an eyes-open paradigm.

It is well-established that brain states are associated with neural oscillations (Singer, 1999) in distinct EEG frequency bands (Klimesch, 1999) linking sustained neurocognitive states to ongoing neural mechanisms (Başar et al., 2001). Our findings recapitulate the dominant finding that alpha power is increased during meditation (Cahn & Polich, 2006; Katyal & Goldin, 2021; Lee et al., 2018; Lomas et al., 2015). Theta on the other hand was not. A recent call for a greater level of spectral and spatial detail in meditation EEG studies (Travis, 2020) highlights that alpha band power is likely constituted by multiple cognitive and cortical processes (Kerr et al., 2011), which may be differentiable by investigating subdivisions of the 7-13 Hz frequency band. Using individual alpha peak frequencies (Klimesch, 1999) aligns potential narrow-band markers, enabling them to be compared across participants, and improves signal to noise. Our approach revealed that amplitudes ranging from the alpha peak frequency (IAF) to 2 Hz above were the most affected.

As alpha findings are not uniform across EEG meditation studies, it is possible that failures to account for individualized alpha could help to explain previous inconsistencies or null findings. Our data and methods allow us to directly test this methodological claim. Using the same statistical models altered only by the frequency binning anchor of the underlying PSD data, our Supplementary Figure 2 analysis demonstrates key elements of spectral “smearing” in fixed-frequency analyses. In our sample, individual alpha peak frequency varied from 8.0 to 12.0 Hz; a 4.0 Hz spread, twice the width of the 2 Hz bands used. Thus, a band defined at fixed frequencies cannot be anatomically or functionally equivalent across participants. As a result, the alpha-peak amplitude measure is weakened under fixed anchoring, whose mean frequency bin of 10.5 Hz notably only matched the true alpha peak of 3/20 individual participants. Smoothing-related differences were not limited to statistical power: with fixed bands, a significant Mindfulness effect appeared in a band 2 - 4Hz below the mean IAF that individualized bands did not support. In data-driven cluster permutation testing, analyses revealed a similar pattern of improved signal to noise under IAF-alignment: aligning spectra to each participant’s peak yielded greater sensitivity to mindfulness-related effects, as well as a larger cluster of discernable spectral difference.

Inconsistent findings in previous mindfulness studies may also reflect the varied designs and testing environments employed. As such, improved standardization and controls have been proposed (Fox et al., 2016; Lee et al., 2018; Matko & Sedlmeier, 2019; Sezer et al., 2022; K. S. Young et al., 2018). Various experimental conditions, including the presence of audiovisual stimulation, can profoundly impact EEG-measured brain signal (Mediano et al., 2024). Alpha band responses to audio and visual content (Mediano et al., 2024) highlight the need to control for detailed aspects of the testing environment manipulation as independent variables. We are not aware of any previous meditation studies that directly investigated the auditory and visual environment (such as natural scenes or auditory instructions being on or off), in a systematically manipulated manner using a within-subjects design.

In our effort to address key stimulus confounds, we observed significant increases in central theta power during auditory instructions across the Mindfulness and Mind-wandering conditions, highlighting the need for detailed study designs that quantify stimulus-related changes. Important for our meditation-related claims, instruction-related theta effects did not overlap spectrally with our alpha bands, and did not interact with Mindfulness. Thus, while posterior alpha increases during Mindfulness were not explained by differences in exposure to auditory instructions, future studies should control stimulus presence across conditions in both design and analysis, to avoid artificially induced spectral effects or masking of true effects. Because auditory and visual stimulation may each modulate alpha-band activity independently of task state (Mediano et al., 2024), periods of guided speech are not neurally equivalent to the silent periods during which participants are actively sustaining the meditative state. Analyses that isolate silent periods therefore may provide the most direct measure of the meditative state itself, isolating the portions of the manipulation that are not influenced by auditory stimuli.

We characterized the effects of naturalistic videos on a mindfulness manipulation to understand the extent to which nature exposure may support early meditation practice (Bell et al., 2026; Djernis et al., 2019; Lymeus et al., 2017). Conceptually, natural environments activate psychological mechanisms of self-transcendence (Anderson et al., 2018), affect regulation (McMahan & Estes, 2015; Richardson et al., 2016), attention (Bell et al., 2026; Berto, 2005), and decreased anxiety (Bratman et al., 2021), all of which are hallmarks of meditation. However, the only observed effect of our video was a Mindfulness × Video interaction in the delta band, which was not part of the primary hypothesis and should be interpreted cautiously. It is possible that our video manipulation simply was not strong enough to introduce a measurable effect on EEG spectral power, or that controlled naturalistic stimuli are insufficient substitutes for “true” natural scenes in facilitating meditation. In any case, the EEG impacts of meditation practice in nature and nature-like environments require future research. The current study provides a roadmap for other systematic manipulations of visual features that may support such manipulations.

Using naturalistic videos (Nute & Chen, 2018) also required our mindfulness manipulation to take place using open eyes. Although many EEG meditation studies have used eyes-closed designs, eyes-open designs are also well established (Lieberman et al., 2025) and allow studying visual effects. Critically, the eyes-open design rules out the possibility that alpha increases during mindfulness are driven by the effect of closed-eyes, a known confound in which closing the eyes is known to increase alpha power (Barry et al., 2007; Ben-Simon et al., 2008). Our findings thus provide useful support to the interpretation that alpha increases during mindfulness practice reflect modulated attention, rather than passive rhythms of the visual system.

There were limitations to our study. Our population was a small convenience-sample of university students, so results may not generalize to more diverse populations. The small sample also limits our ability to characterize individual differences, and we did not measure self-reported attention or mindfulness, or measures of meditation expertise that could link subjective experience and individualized brain responses. Future studies could integrate dynamic elements of self-report [i.e., a neurophenomenological approach] (Berkovich-Ohana et al., 2013; Lutz et al., 2025; Varela & Ura, 1996). Furthermore, meditation effects on neural responses likely vary according to meditation experience (Thomas et al., 2014), depth of meditation attained (Katyal & Goldin, 2021), and specific method of meditation (Lee et al., 2018). Mindfulness is distinguished from other forms of meditation that emphasize repetition (e.g., mantras), reduced cognitive control (e.g., nondirective practice), or compassion-based practices which engage affective and social-cognitive processes (Cahn & Polich, 2006; Fox et al., 2016; Lee et al., 2018). Therefore, the meditative states induced in our manipulation may not be comparable to deeper states of mindfulness or other meditation practices.

Nevertheless, the guided attentional modulation used in this study is representative of light-touch meditation practices that could be used as a potential therapeutic intervention. Mindfulness is often heralded as an effective stress-regulation strategy for non-experts (Borchardt & Zoccola, 2018), and research involving novice meditators may uncover new strategies to ease the adoption of meditation practice such as the adjunctive use of natural scenes or biofeedback strategies (Brandmeyer & Delorme, 2020; Navarro Gil et al., 2018). Further, our manipulation is also ecologically valid given real-world data establishing a mean meditation time of 11 minutes (Cearns & Clark, 2023), and findings that even 5-minutes of mindfulness may combat depression and anxiety in meditation-naive participants (Strohmaier et al., 2021). Short periods of meditation may be a practical target with high translational benefit. Our use of four 5-minute meditation sessions, even punctuated by auditory instructions, ensures that the EEG data collected was sufficient to yield stable spectral power estimates.

Our study is also limited in its methodology. While scalp-recorded power spectral density analyses of EEG rely upon the coordinated activation of large portions of cortex to produce oscillatory impacts detectable at the scalp, other methodologies (e.g., fMRI and invasive electrophysiology) could offer complementary findings to our current study or clarify interpretations (Bauer et al., 2022; K. S. Young et al., 2018). We also did not employ analytical methods such as coherence between electrodes, phase-based analyses, or entropic measures of signal complexity, which may be affected by meditation (J. H. Young et al., 2021) and natural imagery (Bell et al., 2026). Finally, source modelling approaches could be used to estimate the brain regions producing oscillation differences. These methods present further opportunities for analysis.

In summary, guided mindfulness increased posterior upper alpha power, posterior lower alpha 2 power, and amplitude at individuals’ alpha peak frequency relative to guided Mind-wandering. Auditory instructions were associated with increased central theta power, highlighting the importance of modeling task-structure variables in EEG studies of guided meditation. The use of auditory stimuli should be carefully planned and their impacts on neural data should be reported. We did not find evidence that naturalistic video imagery increased EEG markers of meditation. These findings provide additional specificity regarding the alpha-band oscillations associated with mindfulness in novice meditators, and underscore the importance of accounting for auditory task instructions in future meditation EEG studies.

## Data Availability Statement

Data supporting the findings of this study are available from the corresponding author upon reasonable request.

## Author Contributions

DAB – Data curation, Formal analysis, Software, Visualization, Writing – original draft.

KES – Investigation, Project administration, Data curation.

PS – Investigation.

CP – Investigation.

KN – Conceptualization, Resources, Funding acquisition.

NCS – Conceptualization, Supervision, Resources, Formal analysis, Writing – review & editing, Funding acquisition.

CK – Conceptualization, Supervision, Data curation, Formal analysis, Writing – review & editing, Funding acquisition.

---

**Supplementary Figure 1.**
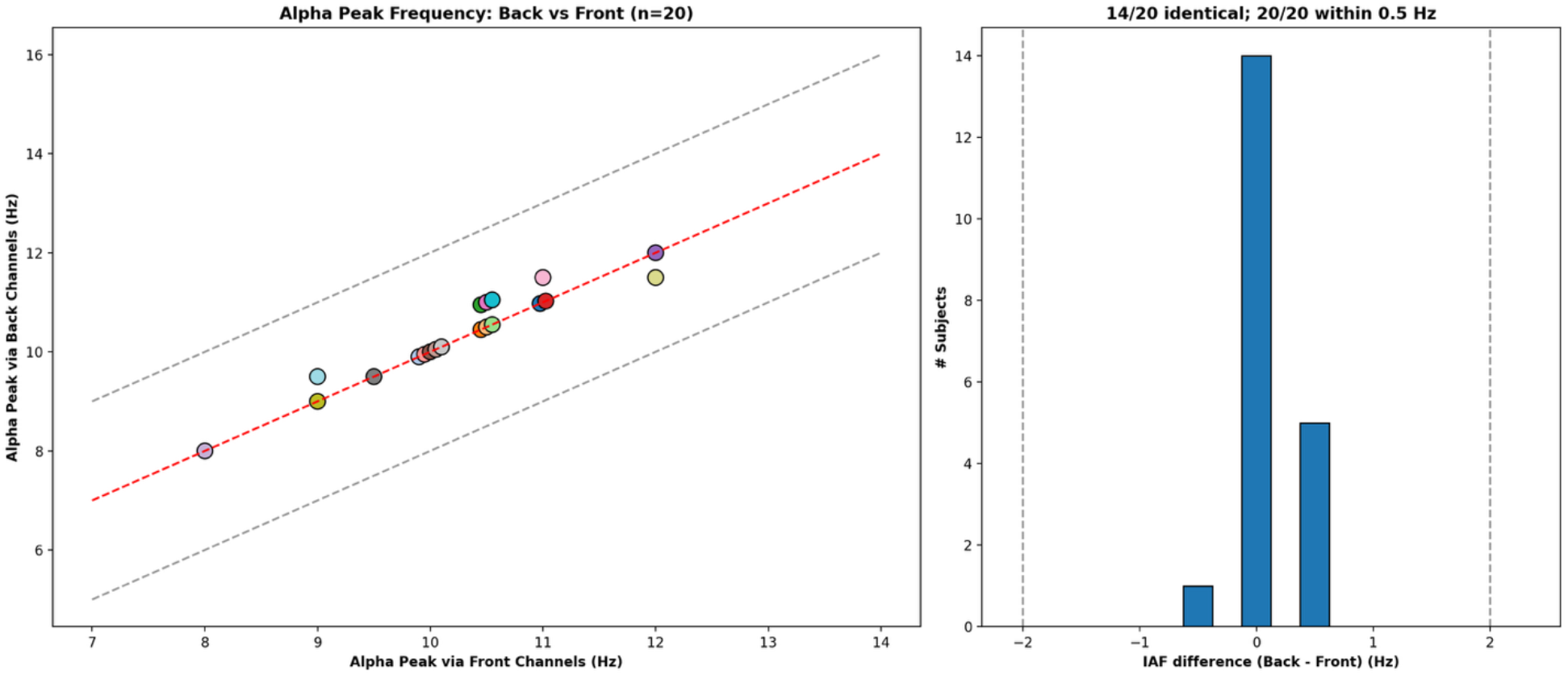
Repeated IAF extraction using a frontal ROI. To ensure posterior channel-derived IAF estimates were consistent across the scalp, IAF derived at posterior and frontal channels were compared. Colored dots (left) show individual participants. Fourteen of 20 subjects showed identical peak frequencies between frontal and posterior channels, and in the remaining six, frontal peaks differed by <= 0.5 Hz from posterior estimates (right). Differences are well below our 2-Hz linear mixed effects analysis bin size, and thus unlikely to influence results.

**Supplementary Figure 2.**
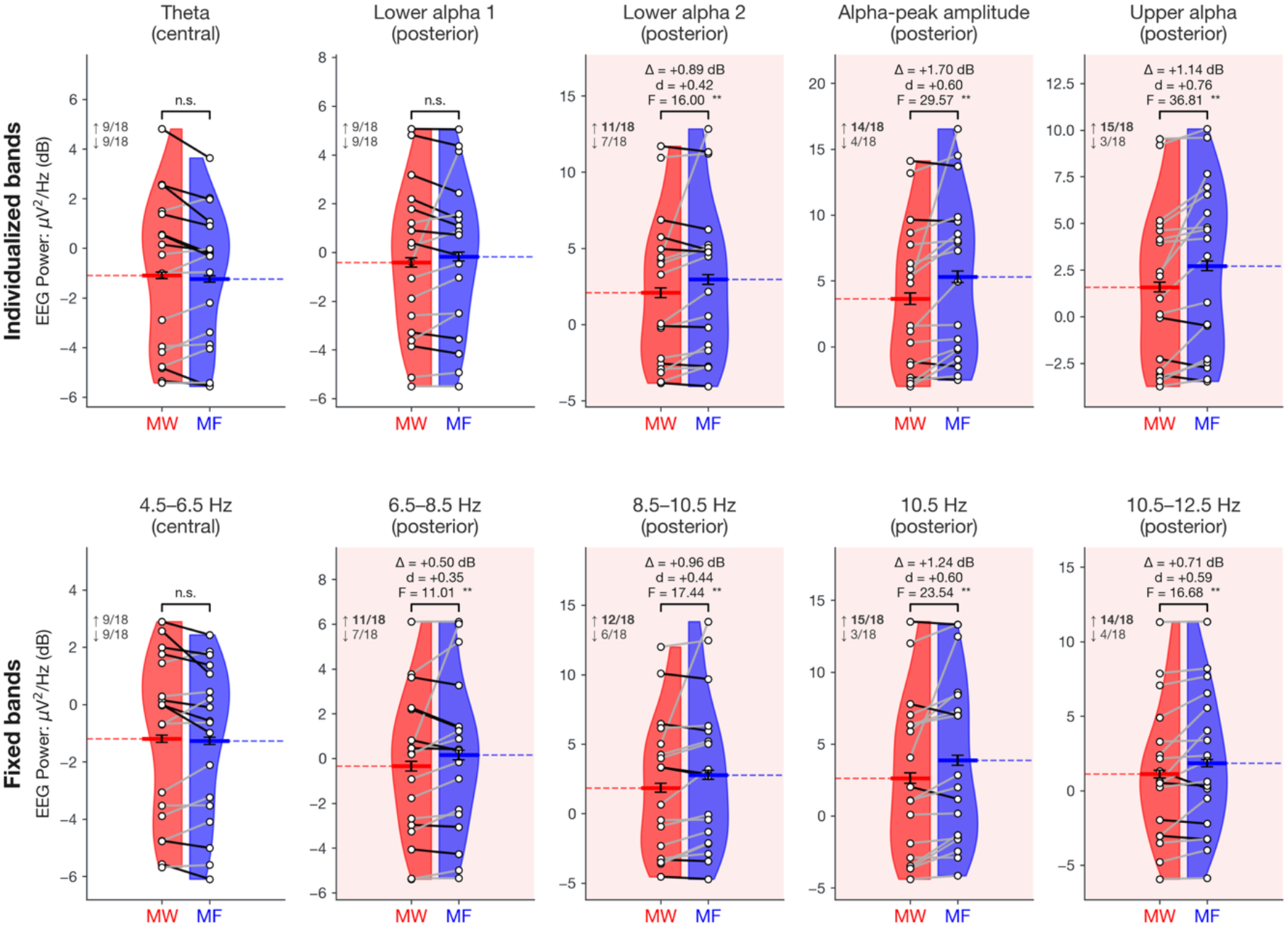
Individualized alpha frequency band alignment concentrates hypothesis-driven findings and increases statistical power. Columns show the five hypothesis-driven spectral outcomes for the Mindfulness contrast. Top row: Band edges are anchored to each participant’s individual alpha peak frequency, as in the main analysis (Figure 4, Table 1). Bottom row: Bands of identical width are instead anchored to the group-mean IAF (10.38 Hz, rounded to the nearest 0.5 Hz bin: 10.5 Hz), producing a non-IAF-adjusted analysis. Bottom row panel titles give the fixed frequency window that each column compares across all participants. The fixed-band method causes mismatched portions of the alpha curve to be averaged across participants – as an example, only 3 of 20 participants had an individual alpha peak at 10.5 Hz, and for the remaining 17 that column does not correspond to their alpha peak. As expected, the fixed-average approach yields a wider set of significant results and lower-magnitude peak effects, consistent with the “smearing” caused by group averaging. Significance indicators reflect FDR-corrected p-values: * p_FDR_ < .05, ** p_FDR_ < .01.

**Supplementary Table 1.** Estimated means and contrasts for interpreted fixed effects. Estimated marginal means and contrasts underlying the linear mixed-effects models reported in Tables 1–2, for reported effects. Estimated means are expressed in power (dB; 10 × log10 of μV^2^/Hz), and a more negative decibel value corresponds to a smaller positive linear value. Differences reflect model-based differences estimated marginal means between conditions, with positive values indicating higher EEG power or amplitude in the first condition listed. Contrasts are shown for hypothesis-relevant effects. Significance indicators for FDR correction: † p < .10, * p < .05 and ** p < .01. The delta Mindfulness × Video rows are simple effects within an interaction that did not survive FDR correction across the 28 interaction tests (q = .203); they are reported for completeness, are not themselves FDR-corrected, and should be treated as exploratory.

| Outcome (ROI) | Est. Marginal Means (SE) |  | Est. Difference | 95% CI | t(df) | p | p<sup>FDR</sup> | Sig. |
| --- | --- | --- | --- | --- | --- | --- | --- | --- |
|  | **Mindfulness** | **Mind-wandering** |  |  |  |  |  |  |
| Theta (Central) | -1.59 (.69) | -1.41 (.69) | -0.17 | [-0.35, +0.01] | -1.91(119.1) | .058 | .245 |  |
| Lower alpha 1 (Posterior) | -0.39 (.71) | -0.62 (.71) | 0.23 | [-0.04, +0.50] | 1.71(119.3) | .089 | .267 |  |
| Lower alpha 2 (Posterior) | 2.69 (1.14) | 1.80 (1.14) | 0.89 | [+0.45, +1.33] | 4.00 (119.3) | <.001 | <.0063 | ** |
| IAF amplitude (Posterior) | 5.06 (1.32) | 3.36 (1.32) | 1.7 | [+1.08, +2.32] | 5.44 (119.4) | <.001 | <.0063 | ** |
| Upper alpha (Posterior) | 2.60 (1.03) | 1.45 (1.03) | 1.14 | [+0.77, +1.52] | 6.07 (119.2) | <.001 | <.0063 | ** |
|  | **Audio Instructions ON** | **Audio Instructions OFF** |  |  |  |  |  |  |
| Theta (Central) | -1.38 (.69) | -1.62 (.69) | 0.24 | [+0.06, +0.41] | 2.69 (119.0) | .008 | .043 | * |
|  | **Video ON** | **Video OFF** |  |  |  |  |  |  |
| Upper alpha (Posterior) | 1.86 (1.03) | 2.19 (1.03) | -0.33 | [-0.69, +0.03] | -1.80 (119.0) | .075 | .261 |  |
|  | **Mindfulness × Video** |  |  |  |  |  |  |  |
| Delta (All) | Mindfulness (Video ON)<br>3.86 (0.49) | Mind-wandering (Video ON)<br>3.58 (0.49) | 0.27 | [-0.01, +0.55] | 1.95 (119) | .054 | — | — |
|  | Mindfulness (Video OFF)<br>3.58 (0.49) | Mind-wandering (Video OFF)<br>3.86 (0.49) | -0.27 | [-0.57, +0.02] | -1.86 (119) | .065 | — | — |

